# Modeling acute kidney injury in kidney organoids with cisplatin

**DOI:** 10.1101/2019.12.22.886572

**Authors:** Jenny L. M. Digby, Aneta Przepiorski, Alan J. Davidson, Veronika Sander

**Author notes:** These authors share senior authorship (Alan J. Davidson, Veronika Sander).

## Abstract

Acute kidney injury (AKI) remains a major global healthcare problem and there is a need to develop human-based models to study AKI *in vitro*. Towards this goal, we have characterized induced pluripotent stem cell-derived human kidney organoids and their response to cisplatin, a chemotherapeutic drug that induces AKI and preferentially damages the proximal tubule. We found that a single treatment with 50 µM cisplatin induces *HAVCR1* and *CXCL8* expression, DNA damage (γH2AX) and cell death in the organoids in a dose-dependent manner but greatly impairs organoid viability. DNA damage was not specific to the proximal tubule but also affected the distal tubule and interstitial populations. This lack of specificity correlated with low expression of the proximal tubule-specific *SLC22A2/OCT2* transporter for cisplatin. To improve viability, we developed a repeated low-dose regimen of 4x 5 µM cisplatin over 7 days and found this causing less toxicity while still inducing a robust AKI response that included secretion of known AKI biomarkers and inflammatory cytokines. This work validates the use of human kidney organoids to model aspects of AKI *in vitro*, with the potential to identify new AKI biomarkers and develop better therapies.

## INTRODUCTION

Acute kidney injury (AKI) is defined by the rapid decline of kidney function and is commonly caused by nephrotoxic side effects of clinical drugs. AKI is a serious condition that predisposes patients to chronic kidney disease (CKD) and mortality. However, there are no targeted therapies for AKI. This is partly due to a lack of clinically relevant models that are required to gain a better understanding of the pathological mechanisms involved in AKI and for the identification of early, reliable biomarkers to improve diagnostic accuracy of AKI (15, 19).

Cisplatin is a chemotherapeutic DNA cross-linking agent that is used to treat solid tumors. The anti-tumor efficacy of cisplatin is limited by severe nephrotoxicity, particularly affecting the S3 segment of the proximal tubule due to the high expression of drug transporters such as *SLC22A2*/*OCT2* (18). The renal pathology of cisplatin-induced AKI is multifactorial, consisting of inflammation, vascular injury, oxidative stress and direct toxicity via generation of reactive oxygen species (11).

In recent years, kidney organoids differentiated from human pluripotent stem cells (hPSC) have been established to study various types of kidney disorders (5, 8). Kidney organoids were reported to be susceptible to proximal tubule injury caused by nephrotoxins, such as cisplatin (2, 9, 16, 17). However, a full characterization of the effects of cisplatin in this system has been lacking.

We have developed a novel kidney organoid protocol that uses a simple, cost-effective method to generate large numbers of organoids, making it ideally suited for injury modeling and drug development (13). Here, we used these kidney organoids to analyze the effects and specificity of cisplatin-induced AKI. We found that cisplatin induces a dose-dependent upregulation of the proximal tubule injury marker *HAVCR1* and the inflammatory cytokine *CXCL8*, DNA damage and cell death that compromised organoid viability. Cisplatin-induced DNA damage was not specific to the proximal tubule but found in the distal tubules and interstitial cells as well. An analysis of cisplatin transporter genes revealed that *SLC22A2*/*OCT2* was expressed at very low levels, providing a potential explanation for the lack of proximal tubule specificity. We found that repeated low doses of cisplatin improved organoid viability, while still inducing injury marker expression and secretion of AKI biomarkers and inflammatory cytokines.

## MATERIALS AND METHODS

### iPSC maintenance and organoid generation

All work was carried out with the approval of Human Participants Ethics Committees (UAHPEC 8712 and HDEC 17/NTA/204) and biosafety. iPSC maintenance and kidney organoid generation were performed as described (13). All experiments were performed using organoids generated from the MANZ-2 iPSC line (13).

### Cisplatin treatment

Cisplatin was diluted from a 5 mM stock in water. For the single high-dose regimen, a whole organoid assay of ∼500-800 organoids was evenly split into 5 wells of a 6-well ultra-low attachment plate (Corning). Cisplatin was added at 0, 5, 10, 25 or 50 µM to ‘stage II’ medium (13) at day (d) 12. Samples were collected 24h and 48h post-treatment, then analyzed using quantitative (q)PCR and immunohistochemistry. For the repeated low-dose regimen, 5 µM cisplatin were added at d12 and subsequently every other day with the medium change for a total of 4 treatments over 7 days. Samples were collected on d19.

### RNA extraction, cDNA synthesis and qPCR

Organoids were washed in PBS and homogenized in TRIzol. Total RNA was extracted using the GENEzol TriRNA Pure kit (Geneaid). RNA from fetal and adult kidney tissue was purchased from Takara. cDNA was synthesized using qScript cDNA SuperMix (Quanta). qPCR was performed using the PerfeCTa SYBR Green FastMix reagents (Quanta) on a QuantStudio 6 Flex Real-Time PCR machine. Gene expression was calculated using the dCt method using *HPRT1* for normalization. Error bars represent standard deviation of triplicate measurements. All qPCR analyses were performed in organoids derived from two independent iPSC lines (MANZ-2 and RiPS or MANZ-4, (13).

### Histology and Immunohistochemistry

Organoids were fixed in 4% paraformaldehyde/PBS. Paraffin embedding and sectioning was performed as described previously (13). Immunohistochemistry (IHC) was performed using standard procedures including heat-induced antigen retrieval. Hoechst 33342 was used for nuclear staining. Antibodies used were: γH2AX (Thermo Fisher CTE2577S and 14-9865-80), MAFB (Novus NBP1-81342), CDH1 (BD Biosciences 610181), MEIS1/2/3 (Active Motif 39796) and LTL (Vectorshield FL-1321). Cell death was measured by TUNEL assay using the ApopTag® Plus Fluorescein *In Situ* Detection kit (Millipore). Fluorescently stained sections were imaged on a Zeiss LSM710 confocal microscope.

### Quantification and statistics

Quantification of IHC was performed on ≥10 organoids per sample using ImageJ. For quantification of marker co-localization, double-positive cells were counted and normalized to the area of the respective tissue marker. Statistical significance was determined using one-way ANOVA or unpaired t-test in Prism (GraphPad). P-values ≤ 0.05 were considered to be statistically significant. * p-value ≤ 0.05; ** p-value ≤ 0.01; *** p-value ≤ 0.001; **** p-value ≤ 0.0001.

### Cytokine array

≥20 organoids per condition were cultured in a single well of a 24-well ultra-low attachment plate in 500 µL stage II medium ± cisplatin for either 48 h (single dose cisplatin regimen) or for the last 48 h of the repeated low-dose regimen. Conditioned culture media were analyzed using the Proteome Profiler™ Human XL Cytokine Array (R&D Systems). Signals were visualized with enhanced chemiluminescence on a Bio-Rad ChemiDoc™ MP Imaging system. Intensities of the duplicate signals were quantified using ImageJ. qPCR was performed to analyze changes in gene expression on RNA isolated from the same organoids as used for the cytokine array, as well as on organoids from independent assays (MANZ-2 and MANZ-4-derived).

## RESULTS

### Cisplatin induces AKI marker expression, DNA damage and cell death in kidney organoids

Previous studies reported cisplatin treatment of kidney organoids with doses ranging from 5-100 μM leading to induction of injury markers, apoptosis and DNA damage (2, 9, 17). To investigate whether these findings were reproducible on kidney organoids generated with our protocol (13), we added single doses of 5, 10, 25 or 50 μM of cisplatin to the culture medium of day (d) 12 organoids and collected samples 24 and 48 hours (h) post-treatment (**Fig. 1*A***). Following the 24h treatment, expression of AKI marker *HAVCR1* (aka *KIM1*) remained unchanged after exposure to 5-25 μM cisplatin but increased 2.4-fold with 50 μM cisplatin. After 48h, both 25 and 50 μM cisplatin induced *HAVCR1* expression significantly (1.7 and 4.1-fold). A similar dose response was observed for expression of the inflammatory cytokine *CXCL8*. No significant change in expression was measured with 5-25 μM cisplatin, whereas 50 μM cisplatin resulted in a 6.5- and 2.4-fold induction of *CXCL8* 24h and 48h post-treatment, respectively (**Fig. 1*B***). To determine the extent of DNA damage and cell death as well as the spatial distribution of cells affected by cisplatin, we used γH2AX antibody and TUNEL staining on paraffin sections of control and cisplatin-treated organoids. Image analysis revealed that both markers were rare in the nuclei of control organoids but increased with 25 and 50 μM of cisplatin (**Fig. 1*C* *and* *D***). γH2AX^+^ and TUNEL^+^ cells were scattered throughout the organoid sections with no obvious accumulation to tubules. These findings suggest that our kidney organoids respond to cisplatin treatment with induction of AKI markers as well as dose-dependent DNA damage and cell death, consistent with previous data (9, 17).

**Fig. 1.**
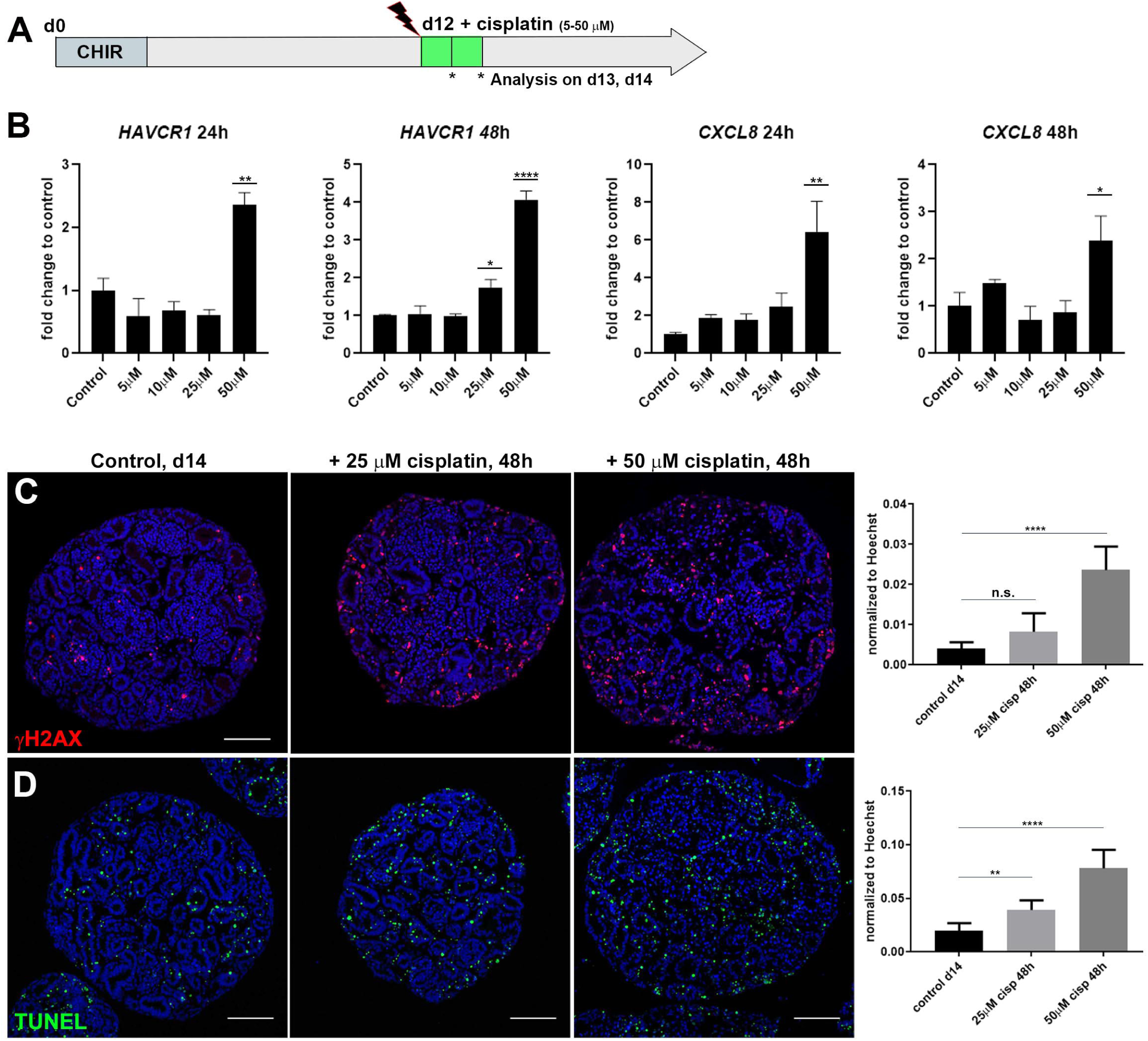
Cisplatin induces tubular injury, DNA damage and cell death in kidney organoids. *A*: Schematic of cisplatin treatment on kidney organoids. *B*: qPCR showing elevated expression of *HAVCR1* and *CXCL8* with increasing doses of cisplatin. *C* and *D*: IHC stainings on paraffin sections of d14 organoids and quantification showing DNA damage marker γH2AX and cell death marker TUNEL increased with 25 and 50 µM cisplatin. n.s., not significant. Scale bars, 100 μm.

### Cisplatin targets interstitial cells

We next investigated if our injury model could recapitulate the proximal tubule-specific cell damage found in AKI patients and animal models. Upon co-immunostaining of γH2AX with antibodies for podocytes, proximal tubule cells, distal tubule cells and interstitial cells, we observed that MAFB^+^ podocytes were the least damaged by cisplatin (**Fig. 2*A***). In contrast, γH2AX co-labeled a subset of LTL^+^ proximal tubule cells, CDH1^+^ distal tubule cells and MEIS1/2/3^+^ interstitial cells in cisplatin-treated organoids (**Fig. 2, *B-D***). Quantification revealed that 50 µM cisplatin induced a 5.5-fold increase in γH2AX^+^ proximal tubule cells and a 3.7-fold increase in distal tubule cells compared to untreated controls (of n≥10 organoids). Strikingly, the largest increase in γH2AX was found in interstitial cells (67-fold; **Fig. 2*E***). This result demonstrates that cisplatin predominantly targets the stromal compartment in kidney organoids.

**Fig. 2.**
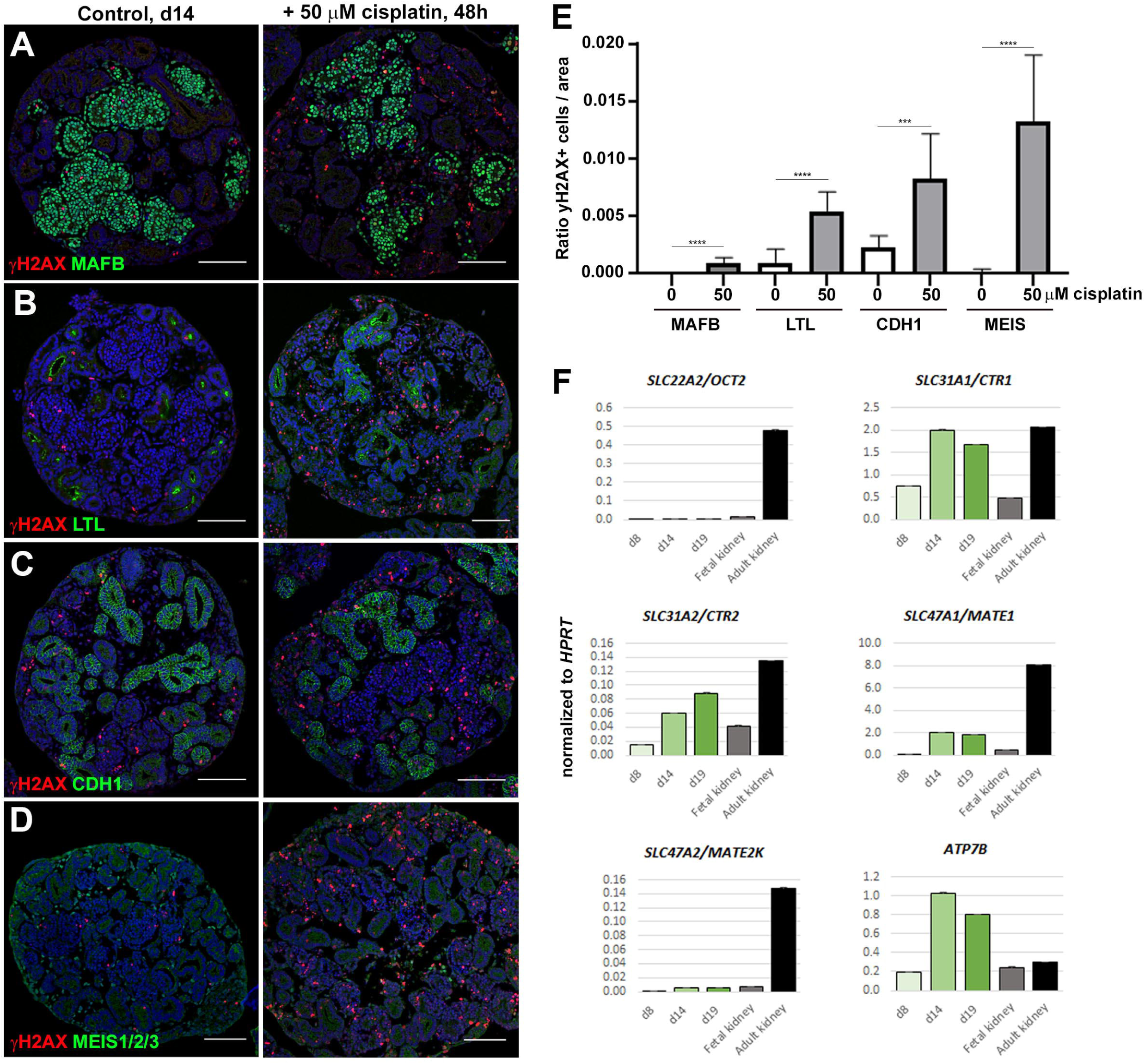
Cisplatin predominantly targets interstitial cells in kidney organoids. *A*-*D*: IHC staining for co-localization of DNA damage marker γH2AX with MAFB^+^ podocytes, LTL^+^ proximal tubules, CDH^+^ distal tubules and MEIS1/2/3^+^ interstitial cells in control and 50 µM cisplatin-treated organoids. *E*: Ratio of γH2AX^+^ cells per marker. *F*: qPCR showing expression of cisplatin transporters in d8, d14 and d19 kidney organoids, and fetal and adult human kidneys. Scale bars, 100 μm.

### Low expression of cisplatin transporters in kidney organoids

To elucidate why tubular cells were less susceptible to cisplatin-induced injury than interstitial cells, we measured the expression levels of cisplatin influx transporters encoded by *SLC22A2/OCT2*, a major cisplatin transporter that is specifically expressed by proximal tubule cells, and *SLC31A1*/*CTR1* and *SLC31A2/CTR2* (4, 10). For cisplatin efflux we measured expression of *ATP7B* and the *SLC47A1*/*MATE1* and *SLC47A2/MATE2K* transporter genes, of which *SLC47A2/MATE2K* is uniquely expressed in the proximal tubule (4, 7). Three stages of organoid development (d8, d14 and d19) were analyzed alongside commercially available RNA of fetal and adult human kidneys using qPCR. We found that expression of *SLC32A1/CTR1, SLC32A2/CTR2, ATP7B* and *SLC47A1*/*MATE1* increased with organoid maturation and was comparable to (or exceeded) the levels of these markers in fetal or adult kidney tissue. In contrast, expression of proximal tubule-specific *SLC22A2/OCT2* and *SLC47A2/MATE2K* transporters was equally low in organoids and fetal kidney (**Fig. 2*F***).

### Repeated low-dose cisplatin treatment reduces cytotoxicity

As shown above, exposure of organoids to 25 and 50 µM cisplatin for 48h leads to acute DNA damage and cell death (**Fig. 1*C*** ***and*** ***D***). At collection on d14, the cisplatin-treated organoids appeared healthy and similar in morphology to untreated controls (not shown). However, treated organoids deteriorated at later stages of culture whereas untreated controls retained a healthy appearance (**Fig. 3*A***). To increase organoid viability and to more closely mimic the repeated dosing regimen of chemotherapy, we tested a repeated low-dose cisplatin regimen on the organoids. To do this, 5 µM cisplatin was added at days 12, 14, 16 and 18 (4x 5 µM) before collection at d19 (**Fig. 3*B***). qPCR revealed a 5.5-fold induction of *HAVCR1* and 19-fold induction of *CXCL8* in 4x 5 µM cisplatin-treated organoids compared to controls (**Fig. 3*C***). Importantly, no tissue disintegration was observed at later stages, indicative of improved organoid viability (**Fig. 3*D***).

**Fig. 3.**
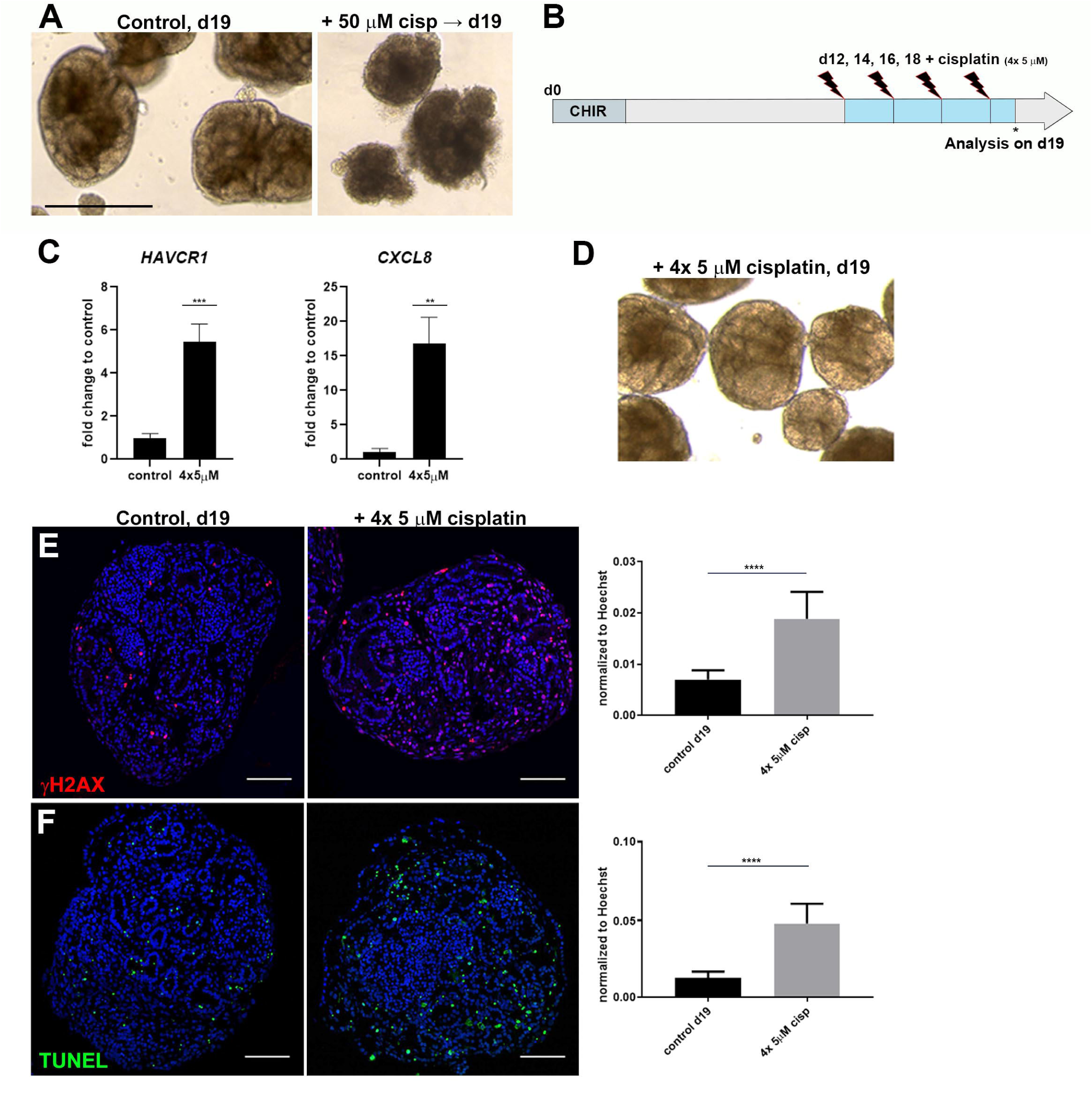
Repeated exposure to low-dose cisplatin reduces cell death and structural deterioration. *A*: Brightfield imaging showing healthy organoids at d19 and organoid deterioration after 50 µM cisplatin. *B*: Schematic of repeated low-dose cisplatin treatment. *C*: qPCR analysis of d19 organoids showing *HAVCR1* and *CXCL8* expression increased upon 4x 5 µM cisplatin. *D*: Organoids treated with 4x 5 µM cisplatin maintain tubular structures. *E* and *F*: IHC showing DNA damage marker γH2AX and cell death marker TUNEL increased with 4x 5 µM cisplatin. Scale bars, 400 μm.

An analysis of 4x 5 µM-treated organoids for γH2AX and TUNEL by IHC showed that both markers increased to a similar extent as with the single treatment (2.7- and 3.6-fold, respectively; **Fig. 3, *D and E***). Co-labeling of γH2AX^+^ nuclei with the tissue markers MAFB, LTL, CDH1, MEIS1/2/3 (**Fig. 4, *A-D***) showed that cisplatin had the greatest effect on interstitial cells (20-fold), whereas for the tubular compartment there was a trend for more proximal tubule injury (3.1-fold) relative to the distal (1.7-fold; **Fig. 4, *E***). Taken together, the repeated low-dose cisplatin treatment recapitulated the induction of kidney injury marker expression observed upon exposure to single high-dose cisplatin, yet exhibited less DNA damage in interstitial cells and improved organoid viability.

**Fig. 4.**
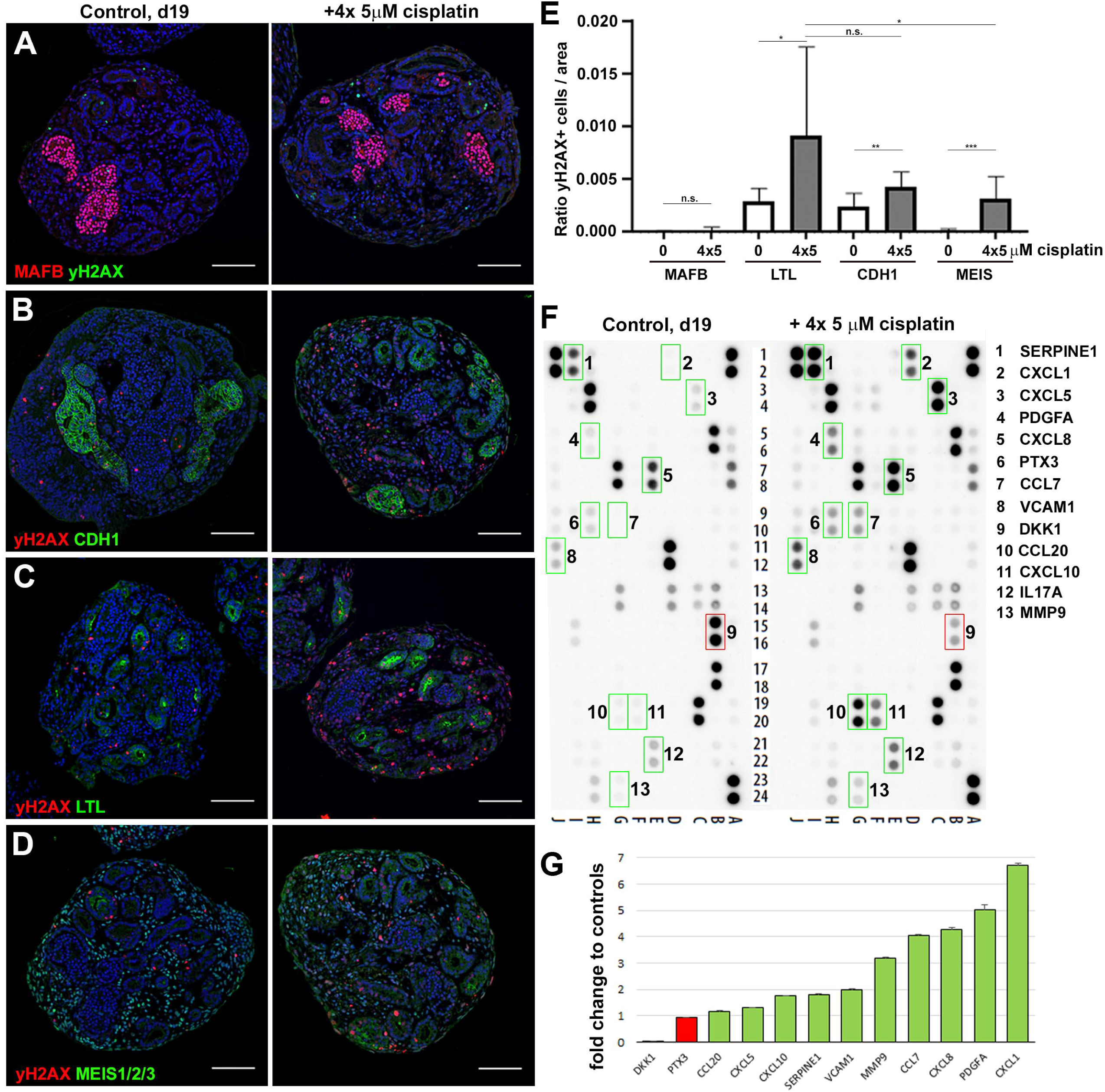
Organoids secrete AKI biomarkers and cytokines in response to cisplatin. *A*-*D*: IHC on d19 control and 4x 5 µM cisplatin-treated organoids for co-localization of DNA damage (γH2AX) with kidney tissues (MAFB^+^ podocytes, LTL^+^ proximal tubules, CDH1^+^ distal tubules, MEIS1/2/3^+^ interstitial cells). *E*: Quantification of γH2AX co-localization with kidney tissues. *F*: Cytokine array analysis using culture media collected from control and 4x 5 µM cisplatin-treated organoids. Factors with higher (green boxes) or lower (red box) secretion in cisplatin-treated versus control organoids are indicated. *G*: qPCR showing transcriptional changes of the cytokines. Expression of *IL17A* was below the detection limit. Scale bars, 100 μm.

### 4x 5 µM cisplatin-treated organoids secrete inflammatory cytokines and AKI biomarkers

Having improved the injury regimen, we next surveyed the cytokines secreted into the medium in response to cisplatin using the Proteome Profiler Human XL Cytokine Array, a membrane-based sandwich immunoassay that can detect 102 cytokines. We found 12 secreted factors with a ≥2-fold increase in signal intensity upon cisplatin treatment (SERPINE1/PAI-1, CXCL1, CXCL5, PDGFA, CXCL8, PTX3, CCL7, VCAM1, CCL20, CXCL10, IL17A and MMP9 (**Fig. 4*F***, green boxes)), and decreased secretion of the Wnt-inhibitor DKK1 (**Fig. 4*F***, red box). qPCR analysis of the genes encoding these factors revealed that differential protein secretion correlated to changes in gene expression (**Fig. 4*G***). As several of these cytokines have been implicated in acute or chronic kidney injury (CXCL1, CXCL5, CCL7, CCL20, IL17A) and renal fibrosis (SERPINE1, PDGFA, MMP9), these data demonstrate the utility of cisplatin-treated kidney organoids as a platform to identify AKI biomarkers and cytokines that model the inflammatory profile seen in AKI (6).

## DISCUSSION

The main finding of this work is that cisplatin, both the single high-dose and repeated low-dose regimen, damages all cellular compartments in the organoids but has the most profound effect on interstitial cells. This is in contrast to previous reports, where a single dose of 5 µM cisplatin was suggested to cause DNA damage and apoptosis specifically to proximal tubules (9, 17). Given the low expression of *SLC22A2*/*OCT2* in our kidney organoids, we speculate that cisplatin uptake occurs via the widely-expressed CTR transporters and thus, accounts for the lack of proximal-tubule specific damage (12). As *SLC22A2*/*OCT2* is not normally expressed in the fetal human kidney, and as kidney organoids represent a fetal stage of differentiation (13, 17), the lack of *SLC22A2*/*OCT2* expression is most likely due to immaturity of our organoids. This lack of specificity may not apply to kidney organoids generated by other protocols that can be cultured for longer periods. The high rate of stromal cell proliferation, as compared to the tubular cells, may be responsible for the enriched susceptibility to DNA damage we observe in these cells, given that cisplatin targets dividing cells (18). Together, our findings suggest that cisplatin exerts a general cytotoxic effect on kidney organoids reminiscent of the impact of the drug on tumor cells, rather than acting as a specific proximal tubule nephrotoxin.

We found that there was a trend for more DNA damage in the proximal tubule over the distal tubule when cisplatin was administered with the repeated low-dose regimen. As this is unlikely due to tubule maturation, we speculate that the 4x 5 µM regimen avoids the acute cell death response of the 50 µM dose, and instead leads to a milder yet accumulative injury phenotype that manifests in robust activation of known AKI markers and cytokines associated with AKI. Such a scenario is more representative of chemotherapeutic low-dose cisplatin administered over an extended period of time, whereby sustained inflammation is a contributing factor to the transition from AKI to renal fibrosis and CKD (14). Our 4x 5 µM approach is also consistent with recent work in mice, where repeated low-dose cisplatin regimens were shown to induce inflammatory cytokines and fibrosis markers in contrast to the acute toxicity, high mortality and lack of fibrosis seen with a single high-dose treatment (1, 3, 15).

Lastly, we show that the proteome array allows for detection of proteins secreted in response to a nephrotoxic insult, analogous to measuring serum and urine markers in AKI patients. This assay provides a proof-of-principle for more sensitive detection tools, such as ELISA, to be used to identify new AKI biomarkers and perform drug-screening approaches. In summary, our organoid model of cisplatin-induced AKI shows limited specificity of injury but a strong representation of the inflammatory response to nephrotoxic insults, and thus provides a valuable system for drug and biomarker discovery that are urgently needed to improve the treatment of patients with AKI.

## ACKNOWLEDGEMENTS

We thank G. Chang for critical reading of the manuscript.

Present address of A. Przepiorski: Department of Developmental Biology, University of Pittsburgh, Pittsburgh, PA, USA

## AUTHOR CONTRIBUTIONS

V.S. and A.J.D. conceptualized the study, J.D., A.P. and V.S. performed experiments, J.D. and V.S. analyzed the data, V.S. and A.J.D. wrote the manuscript, V.S. and A.J.D. supervised the study, and A.J.D. acquired funding.

